# Epstein Barr virus epitope/MHC interaction combined with convergent recombination drive selection of diverse T cell receptor α and β repertoires

**DOI:** 10.1101/2020.02.06.938241

**Authors:** Anna Gil, Larisa Kamga, Ramakanth Chirravuri-Venkata, Nuray Aslan, Fransenio Clark, Dario Ghersi, Katherine Luzuriaga, Liisa K. Selin

**Author notes:** Address correspondence to Liisa K. Selin or Katherine Luzuriaga.

## Abstract

Recognition modes of individual T cell receptors (TCR) are well studied, but factors driving the selection of TCR repertoires from primary through persistent human virus infections are less well understood. Using deep sequencing, we demonstrate a high degree of diversity of EBV-specific clonotypes in acute infectious mononucleosis. Only 9% of unique clonotypes detected in AIM persisted into convalescence; the majority (91%) of unique clonotypes detected in AIM were not detected in convalescence and were seeming replaced by equally diverse “de-novo” clonotypes. The persistent clonotypes had a greater probability of being generated than non-persistent due to convergence recombination of multiple nucleotide sequences to encode the same amino acid sequence, as well as the use of shorter CDR3 regions with fewer nucleotide additions (i.e. sequences closer to germline). Moreover, the two most immunodominant HLA-A2-restricted EBV epitopes, BRLF1**_109_** and BMLF1**_280_**, show highly distinct antigen-specific public (i.e. shared between individuals) features. In fact, TCRα CDR3 motifs played a dominant role, while TCRβ played a minimal role, in the selection of TCR repertoire to an immunodominant EBV epitope, BRLF1. This contrasts with the majority of previously reported repertoires, which appear to be selected either on TCRβ CDR3 interactions with peptide/MHC or in combination with TCRα CDR3. Understanding of how TCR/peptide/MHC complex interactions drive repertoire selection can be used to develop optimal strategies for vaccine design or generation of appropriate adoptive immunotherapies for viral infections in transplant settings or for cancer.

**Importance:** Several lines of evidence suggest that TCRα and β repertoires play a role in disease outcomes and treatment strategies during viral infections in transplant patients, and in cancer and autoimmune disease therapy. Our data suggests that it is essential that we understand the basic principles of how to drive optimum repertoires for both TCR chains, α and β. We address this important issue by characterizing the CD8 TCR repertoire to a common persistent human viral infection (EBV), which is controlled by appropriate CD8 T cell responses. The ultimate goal would be to determine if the individuals who are infected asymptomatically develop a different TCR repertoire than those that develop the immunopathology of AIM. Here, we begin by doing an in depth characterization of both CD8 T cell TCRα and β repertoires to two immunodominant EBV epitopes over the course of AIM identifying potential factors that may be driving their selection.

## Introduction

Over 95% of the world’s population is persistently infected with Epstein Barr virus (EBV) by the fourth decade of life. In the 30% of individuals who are EBV serologically negative upon entering college, primary infection can result in the syndrome, acute infectious mononucleosis (AIM); the frequency of reported symptomatic disease has varied from 25-77% of these young adults (1, 2). AIM symptoms can vary greatly in severity from a mild short flu-like illness to a more severe syndrome with sore throat, lymphadenopathy, splenomegaly, hepatomegaly and debilitating fatigue, which may last for months (1, 2). However, primary infection in the majority of individuals occurs in young childhood and is essentially asymptomatic rarely developing into AIM. A rare 5% of the population appear to never acquire infection and remain EBV serologically negative; severe illness requiring hospitalization has been reported in individuals who acquire primary EBV infection late in life (3). A history of AIM has been associated with an increased risk of subsequent multiple sclerosis (MS) (4) or Hodgkin’s lymphoma (5). EBV infection is also associated with Burkitt lymphoma, nasopharyngeal cancer, hairy leukoplakia in individuals with AIDS, and lymphoproliferative malignancies in transplant patients (5, 6). EBV-associated post-transplant lymphoproliferative disorders can be prevented or treated by adoptive transfer of EBV-specific CD8 T cells (6–8). Defective CD8 T cell control of EBV reactivation may also result in the expansion of EBV-infected, autoreactive B cells in MS (9). Improvement of MS has followed infusion of autologous EBV-specific CD8 T cells in some patients, but not others suggesting that there may be qualitative differences in EBV-specific CD8 T cell responses that need to be better understood (4).

Altogether, these data indicate that EBV-specific CD8 T cells are important for viral control (10). The integration of computational biology and structural modeling approaches to identify TCR antigen-specificity groups and TCR features associated with virologic control (11–16) would facilitate our understanding of how EBV-specific CD8 T cells control EBV replication and contribute to the development of a vaccine to prevent or immunotherapies to modify EBV infection (7, 8, 17).

One of the hallmarks of CD8 T cells is epitope-specificity, conferred by the interaction of the T cell receptor (TCR) with virus-derived peptides bound to host MHC (pMHC) (18–21). The TCR is a membrane-bound, heterodimeric protein composed of α and β chains. Each chain arises from rearrangement of variable (V), diversity (D), joining (J) and constant (C) gene segments (22), resulting in a diverse pool of unique TCRα and β clonotypes. Additions or deletions of N-nucleotides at the V(D)J junctions, specifically at the complementarity-determining region 3 (CDR3) and pairing of different TCRα and β segments further enhance the diversity of the TCR repertoire, estimated to range from 10^15^-10^20^ unique potential TCRαβ clonotypes (23, 24). This diversity enables CD8 T cell responses to a myriad of pathogens.

The CD8 TCR repertoire is an important determinant of CD8 T cell-mediated antiviral efficacy or immune-mediated pathology (16, 23, 25–28). Defining the relationships between early and memory CD8 TCR repertoires is important to understanding structural features of the TCR repertoire that govern the selection and persistence of CD8 T cells into memory. Deep sequencing techniques, combined with structural analyses, provide a high throughput and unbiased approach to understanding antigen-specific TCRαβ repertoires. We (29) and others (30–33) have recently reported that TCRαβ repertoires of CD8 T cell responses to common viruses (influenza, cytomegalovirus, hepatitis C virus) are highly diverse and individualized (i.e. “private”) but “public” clonotypes (defined as the same V, J, or CDR3 aa sequences in many individuals) are favored for expansion, likely due to selection for optimal structural interactions (34).

Studies of influenza A virus in mice (35) and SIV in rhesus macaques (36) have shown that the efficiency with which TCRβ sequences are produced via V(D)J recombination is an important determinant of the extent of TCRβ sharing between individuals (35, 37). Shared TCRβ amino acid sequences required fewer nucleotide additions and were encoded by a greater variety of nucleotide sequences (i.e. convergent recombination). Both of these features are characteristics of TCRβ sequences that have the potential to be produced frequently (35–39) and are also observed in many public TCRs (29, 30, 38–41).

To thoroughly evaluate molecular features of TCR that are important for driving repertoire selection over time following EBV infection, we used direct *ex vivo* deep sequencing of both TCR Vα and Vβ regions of CD8 T cells specific to two immunodominant epitopes, BRLF-1_109_ (YVL-BR) and BMLF-1_280_ (GLC-BM), isolated from peripheral blood during primary EBV infection (AIM) and 6 months later in convalescence (CONV). Each TCR repertoire had a high degree of diversity. However, we noted that persistent clonotypes accounted for only 9% of the unique clonotypes, yet they predominated in both the acute and convalescent phases of infection. An interesting corollary of this finding was that 91% of the unique clonotypes expanded in acute infection were not expanded in convalescence, appearing to be replaced in 6 months by an equally diverse set of *de novo* clonotypes. Expanded clonotypes detected in AIM and CONV, were more likely to be generated in part as a result of convergent recombination than non-persistent or *de novo* clonotypes and had distinct public features (meaning they are shared between donors), which varied by the specific epitope.

## Results

### Patient characteristics

Three HLA-A*02:01+ individuals presenting with symptoms of AIM and laboratory studies consistent with primary infection were studied (**Table S1**) at initial clinical presentation (AIM) and 6 months later (CONV). Direct tetramer staining of peripheral blood revealed that 2.1%±0.5 (mean±SEM) and 1.1%±0.3 of CD8 T cells were YVL-BR and GLC-BM-specific, respectively, in AIM and declined to 0.3%±0.2 and 0.3%±0.1, in CONV. Mean blood EBV load was 3.8±0.9 log_10_ in AIM and 2.6±0.7 log_10_ genome copies/10**^6^** B cells in CONV.

### Persistent dominant clonotypes represent a small fraction of unique clonotypes, with TCRα and β repertoire diversity maintained by the development of *de novo* clonotypes

To examine features that drive selection of YVL-BR and GLC-BM-specific TCRs in AIM and CONV, deep sequencing of TCRα and β repertoires was conducted directly *ex vivo* on tetramer-sorted CD8 T cells at both time points (**Fig 1, S1-2, Table S2**). YVL-BR- and GLC-BM-specific CD8 TCR repertoires in AIM demonstrated inter-individual differences, and were highly diverse; the mean (±SEM) number of unique clonotypes (defined as a unique DNA rearrangement), were not significantly different in CONV (**Fig 1**). Each unique TCRα or TCRβ clonotype detected in AIM that was also detected in CONV was defined as a “persistent” clonotype. Clonotypes were regarded as “non-persistent” or “*de novo*” if they were detected only during AIM or CONV, respectively. A high level of TCR diversity was maintained from AIM to CONV; however, the number of overlapping unique clonotypes detected in both AIM and CONV was small (**Fig 1Ai, Bi**). Only a small fraction of TCRα or β unique clonotypes specific to YVL-BR (6.6±2.2%) and GLC-BM (9.1±4.2%) that were present in AIM were maintained in CONV (YVL-BR:8.7±4.9%; GLC-BM:18.5±5.6%). However, they comprised 57.5±26.2% (YVL-BR) or 75.5±12% (GLC-BM) of the total CD8 T cell response when including their frequency (sequence reads) in AIM and 35.8±10.2% (YVL-BR) or 55.8±13.4% (GLC-BM) in CONV (**Fig 1Aii,Bii**). While the clonotypic composition of YVL-BR and GLC-BM-specific CD8 T cells changed over the course of primary infection, dominant TCR clonotypes detected during AIM tended to persist and dominate in CONV. Altogether, these data indicate that persistent clonotypes made up only a small percentage of unique clonotypes but were highly expanded in AIM and CONV. Surprisingly, the vast majority (91%) of unique clonotypes were not detected following AIM and were seemingly replaced with *de novo* clonotypes in CONV.

**Figure 1:**
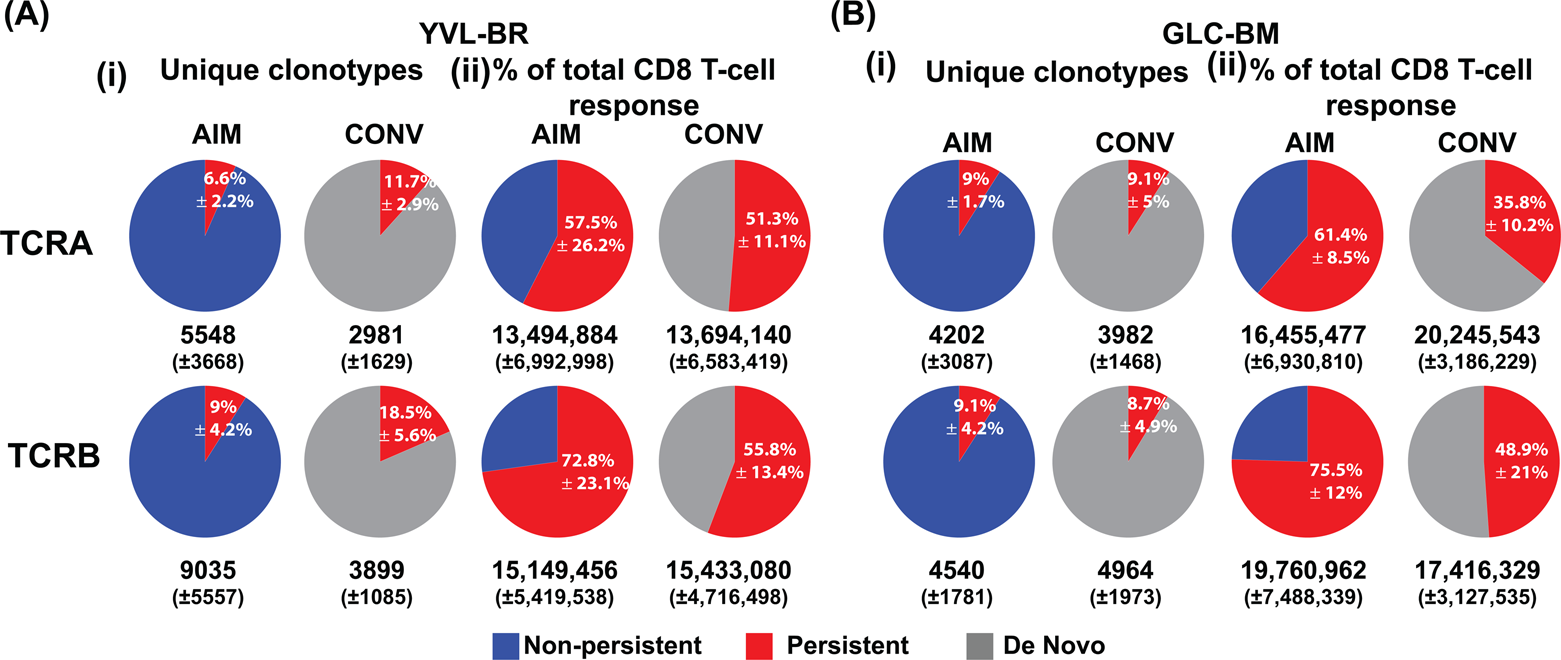
Persistent dominant clonotypes represent a small fraction of unique clonotypes, with TCRα and β repertoire diversity maintained by the development of *de novo* clonotypes (i) Clonotypes that persist from the acute phase into memory represent only 6-18% of the unique clonotypes, but contribute to 35-75% of the total CD8 T cell response. The highly diverse non-persistent clonotypes are replaced by new (*de novo*) highly diverse clonotypes, which were not present in the acute response. The average frequency of unique clonotypes that persist into the memory phase (TRAV and TRBV) in total HLA-A2/YVL-BR-specific (A) and GLC-BM-specific (B) TCR-repertoire is shown (i). The average numbers (±SEM) of unique clonotypes from the 3 donors are shown below the pie charts. Also shown in pie charts is the percentage these clonotypes contribute to the total CD8 T cell response in the HLA-A2/YVL-BR-specific (A) and GLC-BM-specific (B) TCR-repertoire(ii). The average numbers (±SEM) of sequence reads is shown below the pie charts.

### Persistent public clonotypes had an increased probability of generation: convergent recombination contributes to the selection of the persistent TCRα and β repertoire

In both the YVL-BR and GLC-BM TCR repertoires percentage of public clonotypes significantly increased (Chi square: p<0.0001) in the persistent (YVL-BR: TCRAV 34%, TCRBV 17%; GLC-BM: TCRAV 27%, TCRBV 22%) as compared to the non-persistent (YVL-BR: TCRAV 5%, TCRBV 2%; GLC-BM: TCRAV 4%, TCRBV 4%) or de novo repertoires (YVL-BR: TCRAV 5%, TCRBV 1%; GLC-BM: TCRAV 6%, TCRBV 7%). This suggests that the persistent clonotypes may have TCR features that led to greater probability of generation. We tested this by directly calculating the generation probability of amino acid sequences in the CDR3 to determine if the public clonotypes are easier to generate than the private at both time points, acute and convalescent. This allowed a direct and rigorously quantitative test of whether the expanded persistent public clonotypes were of higher generation probability (39, 42). The TCR sequences used by dominant public TCRAV of either GLC-BM and YVL-BR-specific responses have a significantly greater probability of generation while only the GLC-BM TCRBV public but not the YVL-BR public repertoire has a greater probability of being generated (**Fig 1A,B**). This might suggest that TCRAV is dominant and important in the selection of YVL-BR TCR repertoire, while both TCRAV and TCRBV contribute to the GLC-BM TCR repertoire.

To further study this issue we examined whether convergent recombination played a role in the generation of these public persistent TCR (39). Examination of memory antiviral TCRβ repertoires in humans, mice, and macaques suggests that convergent recombination plays an important role in the selection of public antigen-specific TCR (i.e. those shared between individuals of the same haplotype (35–37). Consistent with previous reports for epitope-specific CD8 TCRβ (37, 43, 44) our group found that convergent recombination plays an important role in EBV-specific TCRβ repertoire selection. We also demonstrated that convergent recombination plays a role in selection of persistent TCRα clonotypes specific for the two immunodominant EBV epitopes, YVL-BR and GLC-BM during the course of a human viral infection. There was an increased usage of amino acids derived by multiple different nucleotide (nt) sequences in the CDR3α and β regions of persistent clonotypes as compared to non-persistent and *de novo* clonotypes (**Fig 3A,C**). In fact, we show here that the public TCR had significantly greater usage of these types of amino acids in the CDR3α, as well as the CDR3β (**Fig 3B,D**), as compared to the private clonotypes.

**Figure 2:**
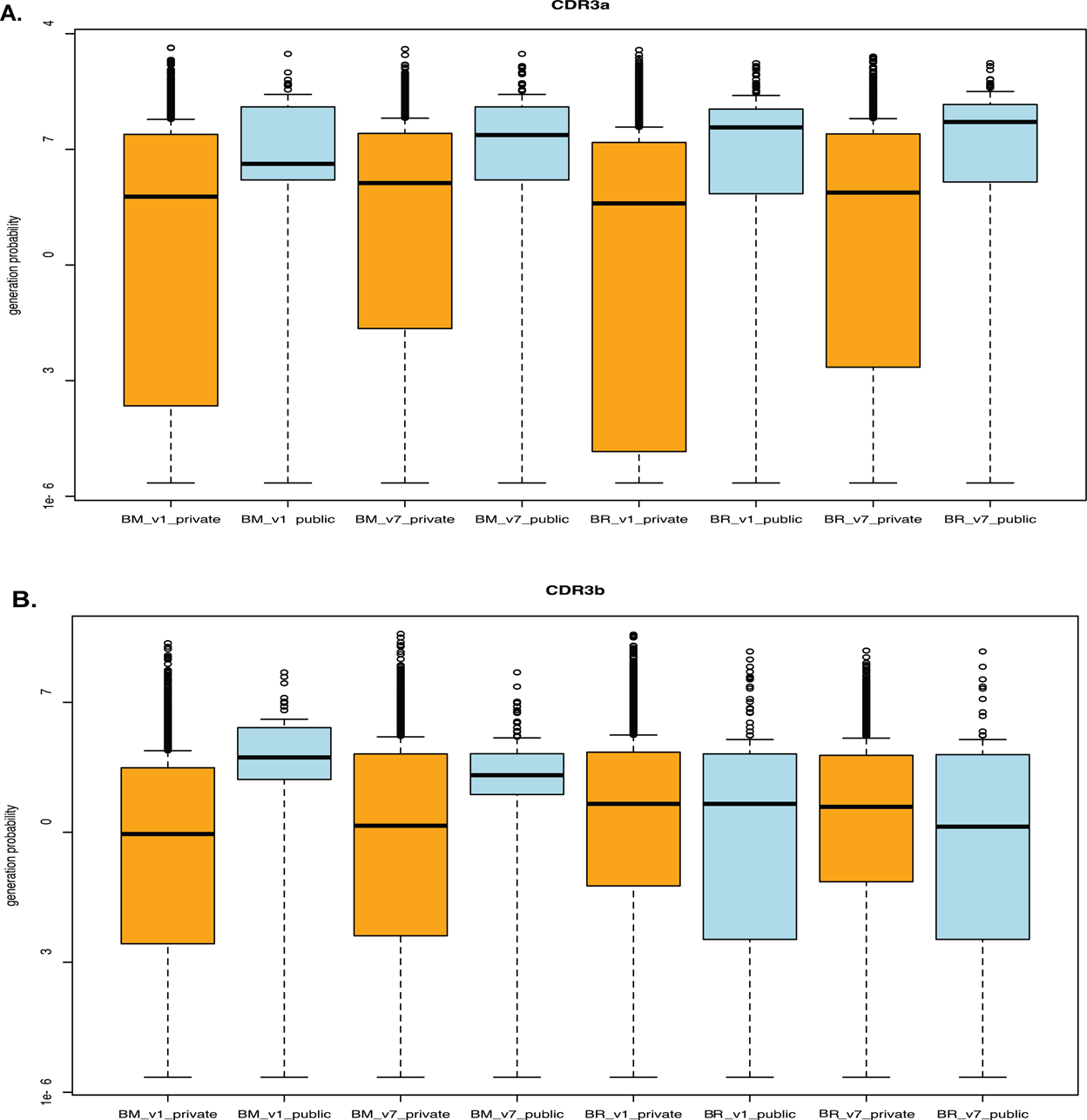
Increased probability of generation of dominant public persistent clonotypes. The algorithm OLGA was used to calculate the generation probability TCR sequences(42). The public TCRAV sequences (A) of either GLC-BM (BM) or YVL-BR-(BR) specific repertoire had a significantly greater probability of generation than private sequences. Only the public TCRBV (B) sequences of GLC-BM but not YVL-BR repertoire had a greater probability of being generated than the private sequences. This is highly consistent with our observation that TCRAV plays a much greater role in the peripheral selection of the YVL-BR TCR repertoire than does TCRBV. The differences between public and private in each pair are all significant (Wilcoxon test, p<0.0001) except TCRBV BR V1 (acute visit 1) and V7 (CONV visit7).

**Figure 3:**
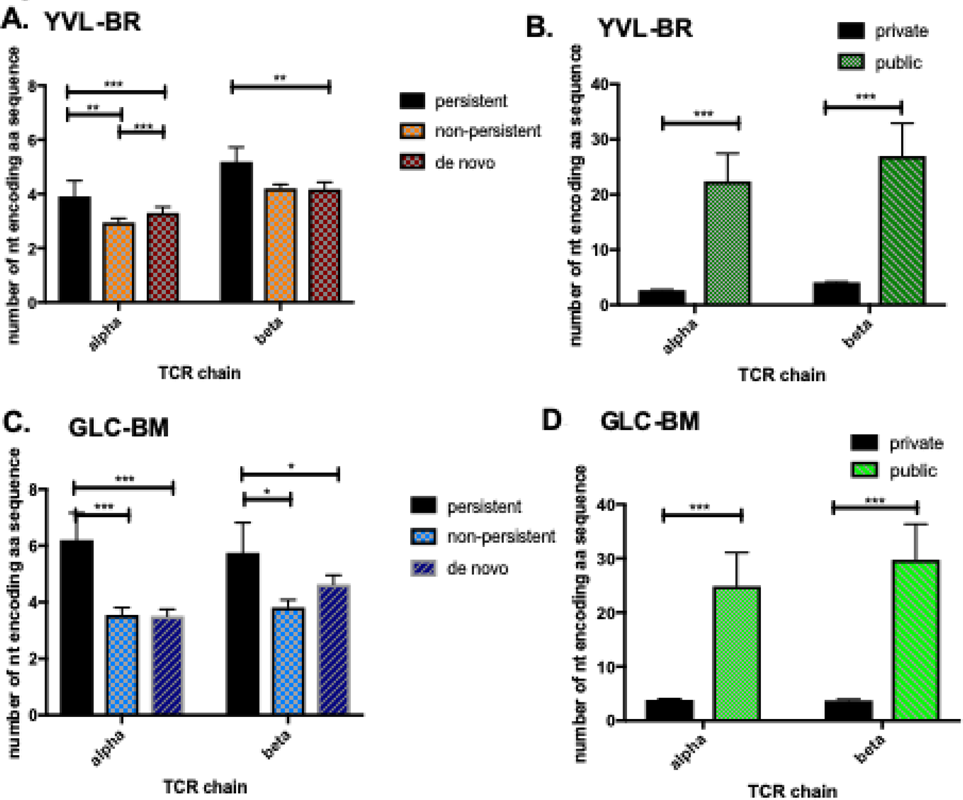
Convergent recombination drives selection of persistent but not non-persistent TCR repertoire: increased usage in CDR3 of amino acids derived by multiple different nucleotide sequences. Number of nucleotides encoding amino acid sequence in CDR3 of YVL-BR-(A,B) and GLC-BM-specific (C,D) TCRα and TCRβ of persistent, non-persistent and *de novo* repertoires (A,C) and private versus public clonotypes (B,D). (A public TCR is defined as more than one donor is using that clonotype based on amino acid sequence). Data was analyzed by two-way ANOVA multivariant analysis with correction for multiple comparisons, * p<0.05, ** p<0.01, *** p<0.001, **** p<0.0001. Error bars are SEM.

Another TCR feature that leads to increased probability of generation is the use of decreased numbers of nucleotide additions in the CDR3, consistent with encoding of the TCR by predominantly germline gene segments (39). This was indeed the case for YVL-BR and GLC-BM-specific clonotypes (**Fig 4**); the CDR3α of persistent YVL-BR- and GLC-BM-specific clonotypes had fewer nucleotide additions compared to non-persistent and increased number of nt additions in *de novo* clonotypes of EBV-BR. However, the CDR3β of persistent YVL-BR- and GLC-BM-specific clonotypes did not have fewer nucleotide additions compared to non-persistent (**Fig 4A,D**). Public clonotypes of each epitope-specific response also had fewer nucleotide additions than private clonotypes except interestingly for YVL-BR CDR3β, where the private had fewer (**Fig 4B,E**). Interestingly, there was an increased usage of glycines in the longer CDR3 of the *de novo* TCR repertoire (**Fig 4C,F**), which has been reported to be a feature associated with greater TCR promiscuity (45, 46). Overall, these results suggest the use of shorter CDR3 regions with fewer nucleotide additions in the persistent TCRAV but not in the TCRBV clonotypes. Curiously, consistent with probability generation data (**Fig 2**) the public TCRBV of EBV-BR were actually significantly longer with increased nucleotide additions.

**Figure 4.**
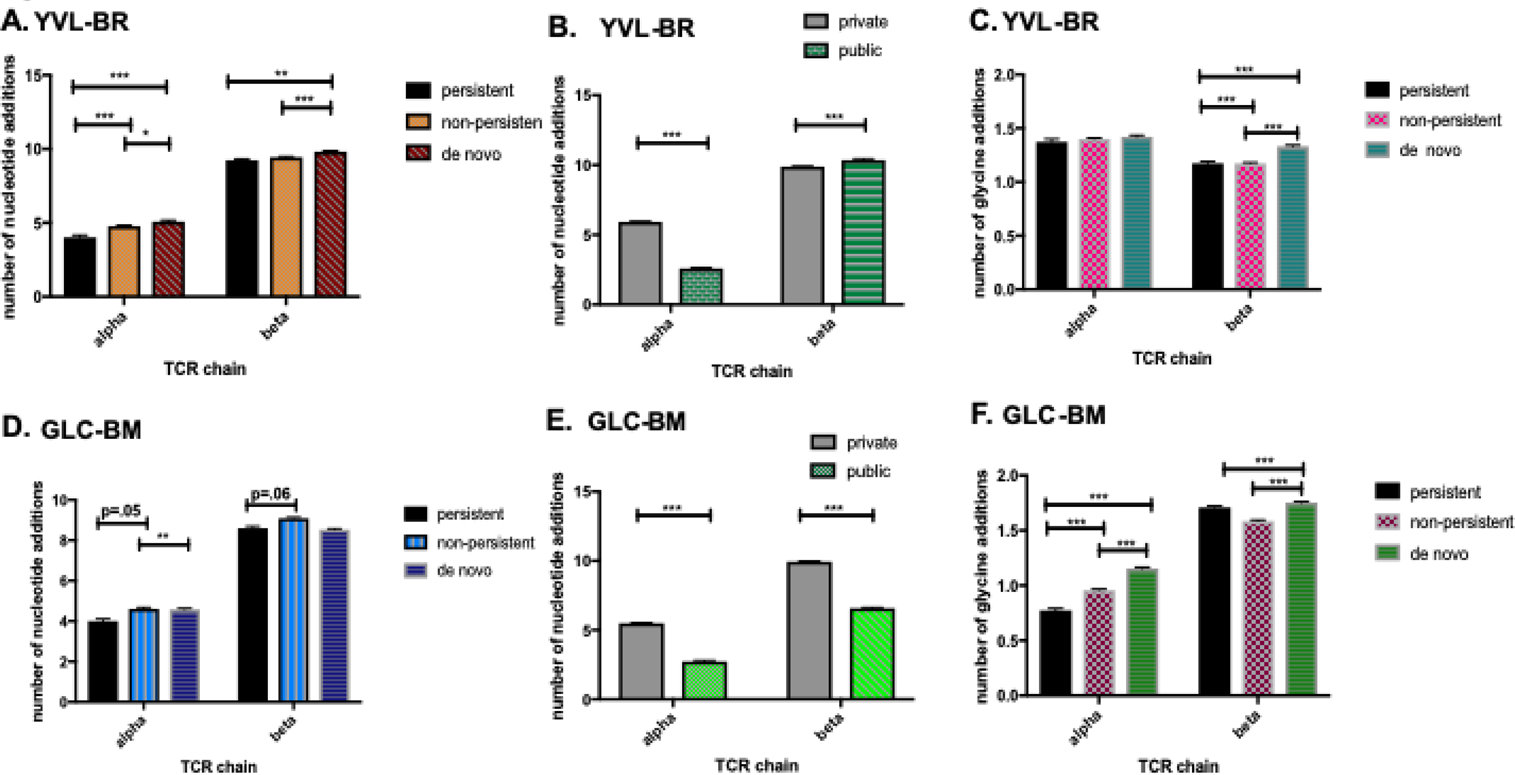
Convergent recombination drives selection of persistent but not non-persistent TCR repertoire: decreased number of nucleotide additions in CDR3 of persistent EBV-specific TCR repertoire. Number of nucleotide additions in the CDR3 of YVL-BR-(A,B) and the GLC-BM-specific (D,E) TCRα and TCRβ of persistent, non-persistent and de novo repertoires (A,C) and private versus public clonotypes (D,E). Increased usage of glycines in the longer CDR3 of the *de novo* TCR repertoire (C,F). Data was analyzed by two-way ANOVA multivariant analysis with correction for multiple comparisons, * p<0.05, ** p<0.01, *** p<0.001, **** p<0.0001. Error bars are SEM.

### CDR3 lengths are a major factor in the selection of the YVL-BR- and GLC-BM-specific TCRα and β repertoires

Differences in dominant YVL-BR- and GLC-BM-specific CDR3α and β lengths were also observed between the epitopes and from AIM to CONV and between persistent and non-persistent or *de novo* clonotypes (**Fig 5**). There were differences in preferential use of CDR3 lengths between YVL-BR and GLC-BM. For instance, the AIM YVL-BR-specific repertoire used more of the shorter 10-mer CDR3β than GLC-BM in both AIM and CONV (**Fig 5Aii**). Within the YVL-BR response, use of the shorter 9-mer CDR3α decreased from AIM to CONV (**Fig 5Ai**). Persistent YVL-BR-specific clonotypes used significantly more of the shorter 9-mer CDR3α and 10-, 11-, and 12-mer CDR3β than the non-persistent. In contrast, the *de novo* clonotypes favored the longer 12-mer CDR3α and focused more on 11-mer CDR3β length (**Fig 5Bi-ii**). Significant changes in the GLC-BM-specific CDR3 length were also observed between AIM and CONV. For example, the frequencies of the longer GLC-BM-specific 12-mer CDR3α and β clonotypes significantly increased from 13.6±6% and 6±2.8%, respectively, in AIM to 24±5% and 17.9±8%, respectively, in CONV, while use of the shorter 11-mer CDR3α decreased (**Fig 5Ai-ii**). The persistent clonotypes preferentially used 9- and 11-mer CDR3α while *de novo* used longer 12- and 14-mer lengths (**Fig 5Biii-iv**). The persistent clonotypes also used 11- and 13-mer CDR3β, while *de novo* used 12-mer lengths.

**Figure 5:**
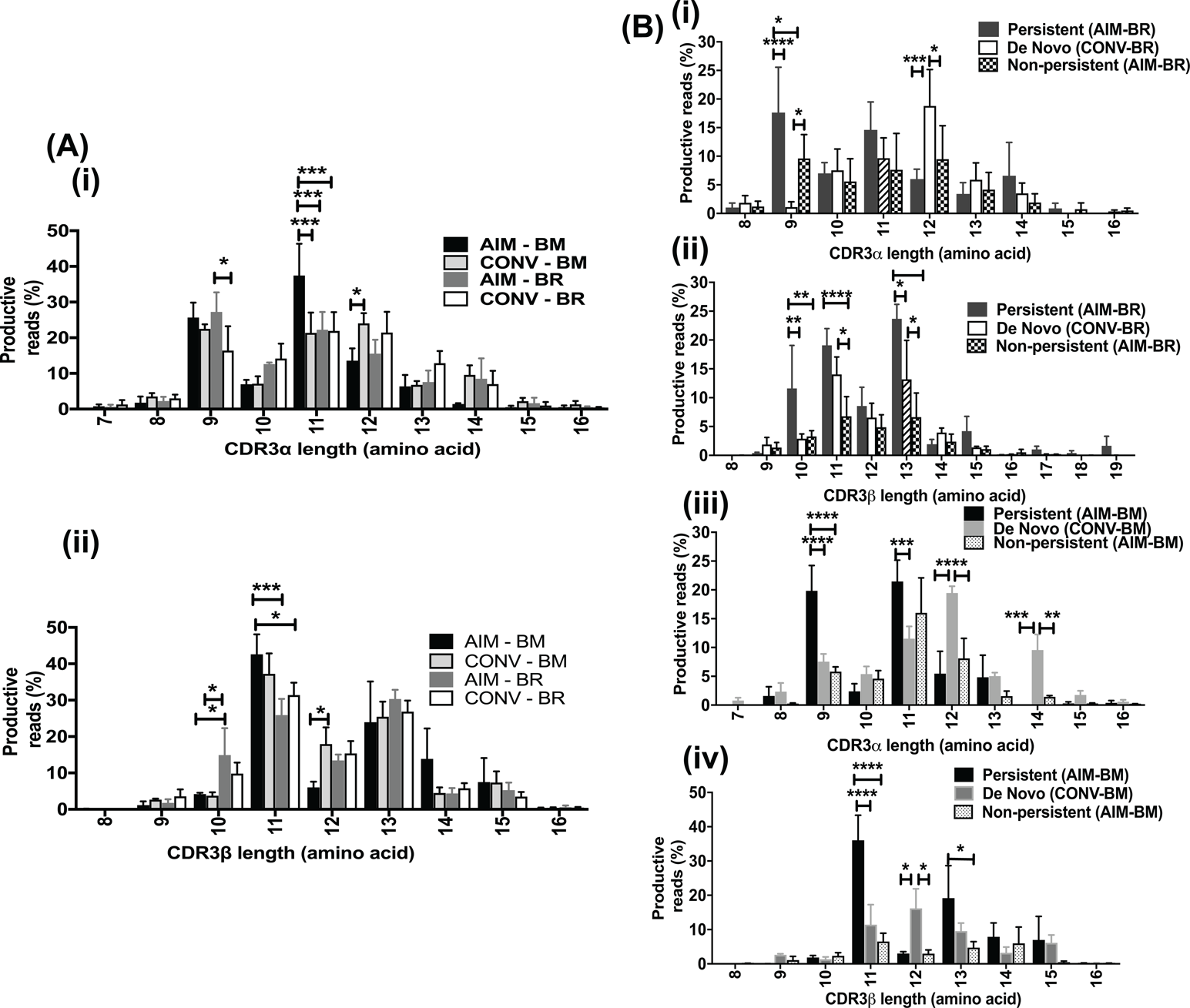
CDR3α and β length distribution of YVL-BR- and GLC-BM-specific CD8 T cells in AIM and CONV (A) and in persistent, *de novo* and non-persistent clonotypes differ (B). (A) The mean CDR3 length distribution of the 3 EBV-infected patients’ TCR repertoire was analyzed by deep-sequencing of tetramer sorted cells during AIM and CONV. (B) The TCR repertoires were analyzed also after dividing each patients samples into 3 groups, those that persist from AIM into CONV and those which do not as well as *de novo* clonotypes arising in CONV. Data was analyzed by two-way ANOVA multivariant analysis with correction for multiple comparisons, * p<0.05, ** p<0.01, *** p<0.001, **** p<0.0001. Error bars are SEM.

### Selection of the TCRα and β repertoires was based on the features on the specific epitope

To further elucidate factors that are driving selection of TCR specific to the two immunodominant EBV epitopes, the characteristics of the TCR repertoires for each of 3 donors were elucidated by systematically analyzing preferential TCRAV or BV segment usage hierarchy as presented in pie charts, CDR3 length analyses, V-J pairing by circos plots of the clonotypes with the dominant CDR3 lengths, and dominant CDR3 motif; the latter determines if there was an enrichment of particular amino acid residues at specific sites potentially important for ligand interaction. Enrichment for certain characteristics would suggest that these features are important for pMHC interaction. (11, 29, 47–50).

### The 9-mer TCR AV8.1-VKDTDK-AJ34 drives selection of YVL-BR-specific CD8 T cells

The YVL-BR-specific TCRα repertoire was focused on one dominant family, AV8, used by all donors in AIM and CONV (**Fig 6Ai, S1Ai**). Similar strong selection bias was not observed in YVL-BR-specific TCRBV usage; there was a great deal of inter-individual variation and preferential usage of multiple families, including BV6, BV20, BV28, BV29 (**Fig 6Bi, S1Bi**). Interestingly, in CONV, some TCRAV and BV gene families that dominated in AIM became extinct or subdominant, or new dominant genes emerged (**Fig 6Ai, Bi**).

Circos plot analyses of the pronounced 9-mer clonotypes showed that the dominant *AV8.1* gene almost exclusively paired with *AJ34* (**Fig 6A, S1Aiii**). CDR3α motif analysis revealed a pronounced motif, “VKDTDK”, in these shorter 9-mer clonotypes, representing 13.8%±5.6 of the total CD8 T cell response during acute AIM (**Fig 6Aiii, S1Aiv, Table S3A**); 87%±1.7 of the clonotypes using this motif were AV8.1 and 92%±1.7 were AJ34. Interestingly, this motif was present in multiple other AV and AJ pairs, including AV12, AV21 and AV3. Obligate pairing of the dominant AV8.1 response to AJ34 containing the highly conserved motif, VKDTDK, was observed in all donors from AIM through CONV, suggesting that the 9-mer AV8.1-VKDTDK-AJ34 expressing clones were highly selected. There was a preferential usage of BV20-BJ2.7 pairing within the dominant 11-mer response (**Fig 6Bii, S1Biii**), without an obvious CDR3β motif (**Fig 6Biii, S1Biv**), highlighting a great degree of diversity in the amino acid sequences. Within the 13-mer response (**Fig 6Biii,S1Biv, Table S3B**), the CDR3β motif, “LLGG”, was commonly used. Clonotypes with this motif were only a minor part of the overall responses in 2 donors (E1603, E1655), but composed 17.4% of the total YVL-BR TCRβ repertoire in E1632.

**Figure 6:**
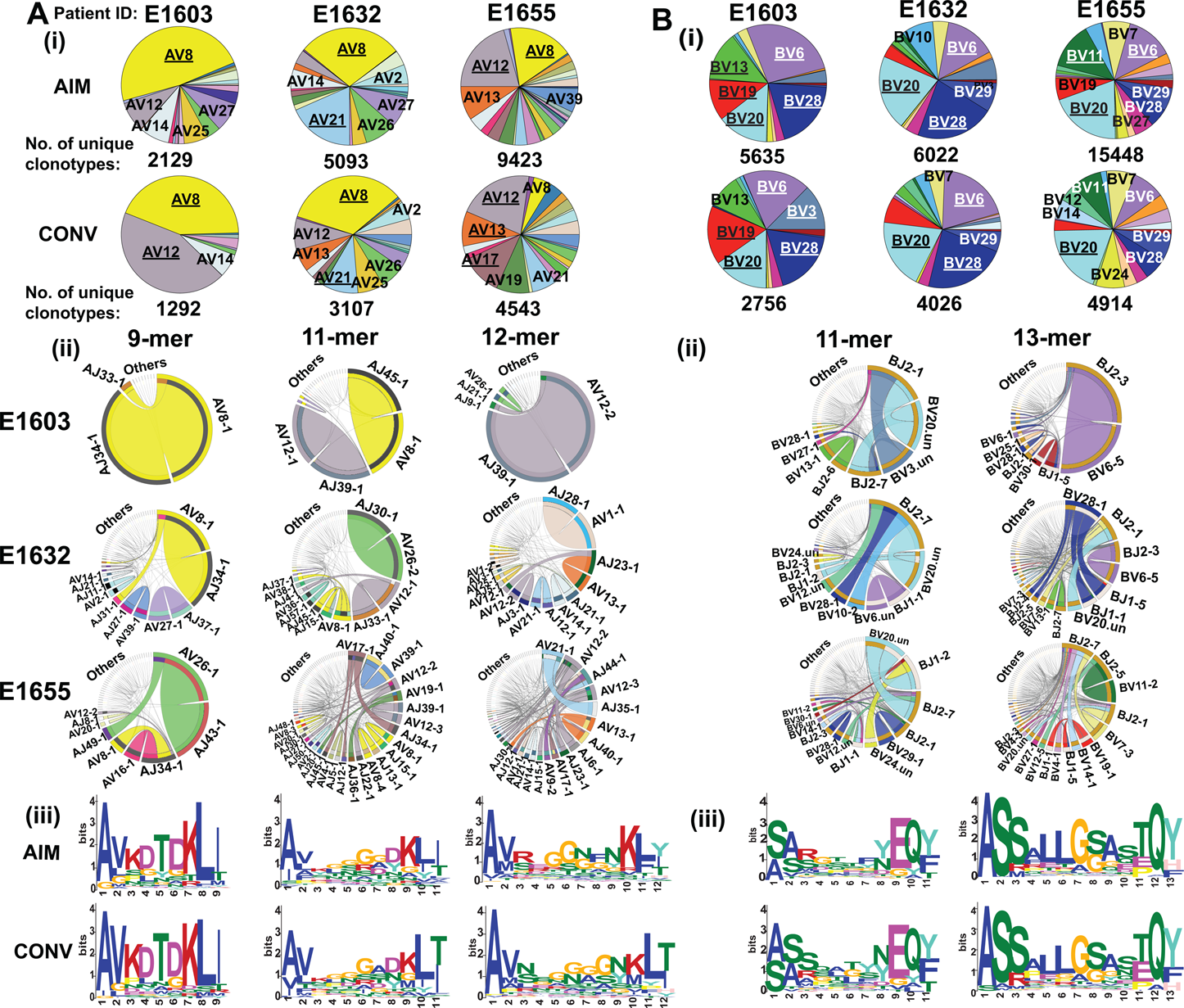
9-mer TCRα AV8.1-VKDTDK-AJ34 drives the selection of YVL-BR-specific CD8 T cells in AIM and CONV. HLA-A2/YVL-BR-specific TCRα (A) and TCRβ (B) repertoires were analyzed for 3 AIM donors (E1603, E1632, E1655) during the acute (within two weeks of onset of symptoms; primary response) and convalescent (6 months later; memory response) phase of EBV infection. Frequency of each TRAV (A) and TRBV (B) in total HLA-A2/YVL-BR-specific TCR-repertoire is shown in pie charts (i). The pie plots are labeled with gene families having a frequency ≥10% (dominant, underlined) or between 5% and 10% (subdominant; not underlined). The total numbers of unique clonotypes in each donor is shown below the pie charts. (ii) Circos plots depicting V-J gene pairing and (iii) CDR3 motif analysis for the clonotypes with the two most dominant CDR3 lengths. Circos plots are only shown for the memory phase (AIM circos plots in Fig 7,8 and S1). The frequencies of V-J combinations are displayed in circos plots, with frequency of each V or J cassette represented by its arc length and that of the V-J cassette combination by the width of the arc. “.un” denotes V families where the exact gene names were unknown.

**Figure 7:**
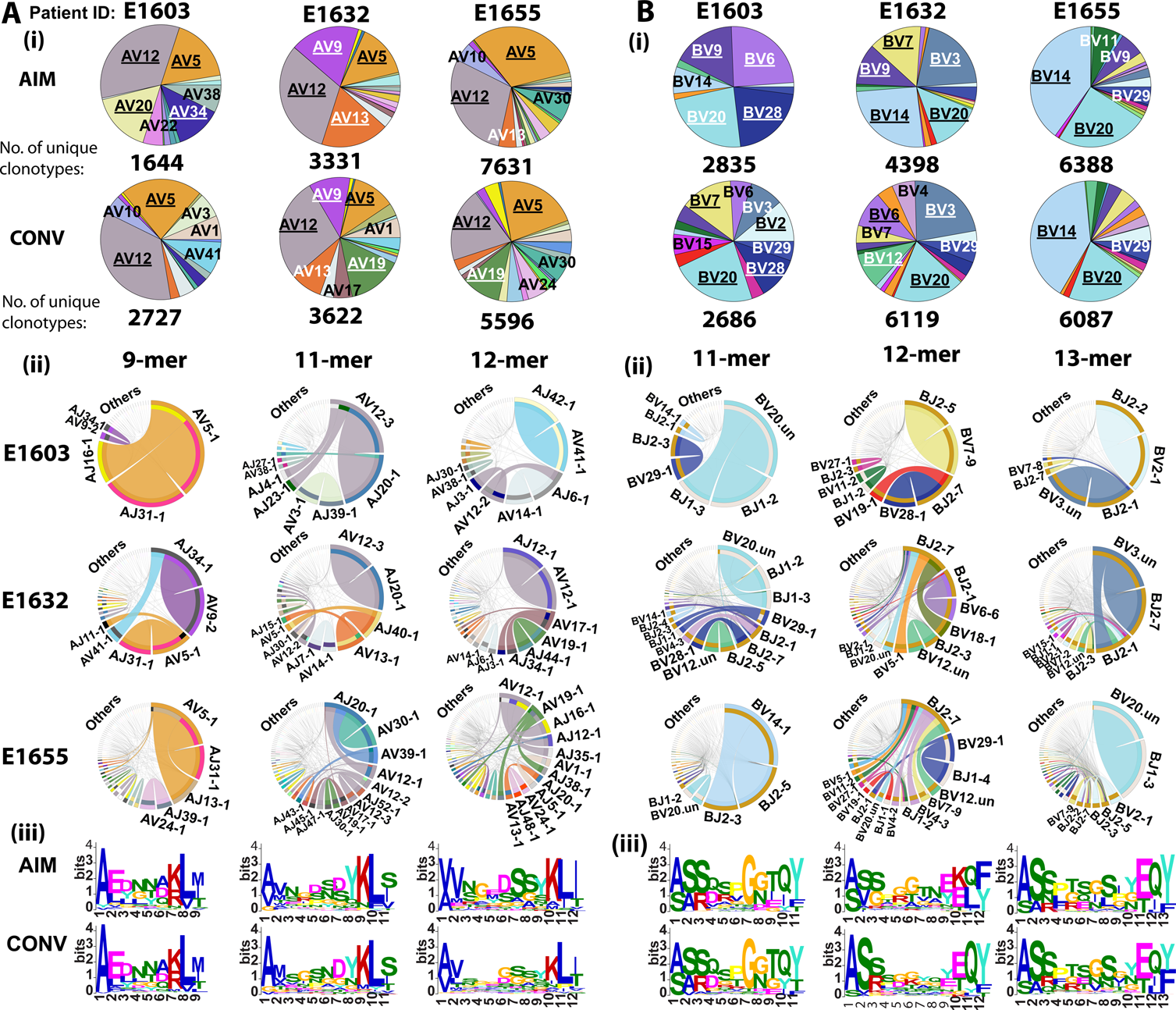
TCRα, AV5-EDNNA-AJ31, and TCRβ, BV14-SQSPGG-BJ2 and BV20-**SARD-BJ1, clones are dominant selection factors for GLC-BM-specific CD8 T cells in AIM and CONV.** HLA-A2/GLC-BM-specific TCRAV (A) and TCRBV (B) repertoires were analyzed for 3 AIM donors (E1603, E1632, E1655) during the acute (within two weeks of onset of symptoms; primary response) and convalescent (6 months later; memory response) phase of EBV infection. Frequency of each TRAV (A) and TRBV (B) in total HLA-A2/GLC-BM-specific TCR-repertoire is shown in pie charts (i). The pie plots are labeled with gene families having a frequency ≥10% (dominant, underlined) or between 5% and 10% (subdominant; not underlined). The total numbers of unique clonotypes in each donor is shown below the pie charts. (ii) Circos plots depicting V-J gene pairing and (iii) CDR3 motif analysis for the clonotypes with the two most dominant CDR3 lengths. Circos plots are only shown for the memory phase. (AIM circos plots in Fig S2-4). “.un” denotes V families where the exact gene names were unknown.

Altogether, these results suggest that the 9-mer AV8.1-VKDTDK-AJ34 expressing clones were highly preferentially selected by YVL-BR ligand during AIM and CONV and that this TCRα could pair with multiple different TCRβ, as suggested by the fact that there was no such dominant TCRβ clonotype. These findings have been independently confirmed using single cell sequencing (51).

### AV5-EDNNA-AJ31, BV14-SQSPGG-BJ2, and BV20-SARD-BJ1 GLC-BM-specific CD8 T cells are highly selected

GLC-BM-specific TCRAV and BV use also had clear preference for particular gene families, maintained from AIM to CONV, consistent with prior reports (52, 53). We observed apparent preferential use of public AV5, 12, and BV20, 14, 9, 28, 29 families (**Fig 7Ai, Bi, S2Ai, Bi**). Like YVL-BR, there were some individual changes in the transition into CONV (**Fig 7Ai,Bi**). Circos plot analysis of the dominant 9-mer CDR3α length clonotypes revealed a conserved and dominant AV5-AJ31 pairing in all 3 donors (**Fig 7Aii, S2Aiii, S3**). A prominent motif, “EDNNA”, was identified within 9-mer clonotypes, of which 85%±11 were associated with AV5-AJ31 (**Fig 7Aiii, S2Aiv, Table S3C**). This CDR3α motif was used by only 2.8%±1.7 of all clonotypes recognizing GLC-BM in the 3 donors. The 11-mer CDR3β BV14-BJ2 pairing exhibited a conserved, previously reported public motif, “SQSPGG” (54), which represented 26% and 40% of the total GLC-BM-specific response in donors E1632 and E1655 in AIM, respectively (**Fig S2Bii-iv, Table S3D**). Within the CDR3β 13-mer response, a conserved BV20-BJ1 pairing, including the previously reported public motif, “SARD”, was used by all 3 donors, and represented 11%±6 of the total GLC-BM-specific response (**Fig 7Biii, S2Bii-iv, Table S3D**). Within the 13-mer CDR3β response, there was also a consensus motif, “SPTSG” present in all 3 donors, which was used by multiple different BV families, which represented 20% and 2% of the total response in donors E1632 and E1655, respectively in AIM (**Fig 7Bii-iv, Table S3D**). These data suggest that, in contrast to YVL-BR, whose TCR-repertoire selection was primarily driven by TCRα, the selection of the GLC-BM-specific TCR-repertoire in AIM was driven by a combination of both TCRα and β.

Overall, despite individual changes, the dominant TCRV gene families and CDR3 motifs that were identified in AIM to drive the selection of YVL-BR or GLC-BM-specific CD8 T cells were predominantly conserved in CONV, suggesting the strength of these TCR features in driving selection of the repertoire (**Fig 6**-**7, Table S3**).

### Persistent, non-persistent, and *de novo* clonotypes differ in selection factors

To address whether clonotypes that persisted into memory show similar characteristics to those that dominate in acute infection, YVL-BR and GLC-BM TCRα/β repertoires were compared between AIM and CONV. The TCR repertoire of persistent and non-persistent clonotypes in AIM, and *de novo* clonotypes in CONV, were examined in order to identify selection factors that governed TCR persistence.

*YVL-BR persistent, non-persistent, and de novo clonotypes have unique characteristics*.

Persistent YVL-BR clonotypes maintained the major selection factors that were identified in AIM (**Fig S3,S4, 8A, Table S4**). Although some features were maintained in all 3 TCR subsets, there were significant structural differences in these repertoires.

The YVL-BR non-persistent CDR3α clonotypes used AV8.1 but it was paired with many more AJ gene families (**Fig S3**). Moreover, AV8.1-VKDTDK-AJ34 clonotypes, which were present in 42±20% or 19±11% of all persistent clonotypes during AIM or CONV, respectively, were present in the non-persistent response at a much lower mean frequency (6±1%; **Fig 8A, Table S4A,B**). The clonal composition of the CDR3β non-persistent response varied greatly in BV family usage between donors (**Table S4D,E**) and lacked identifiable motifs, suggesting that for YVL clones expressing AV8.1-VKDTDK-AJ34 to persist, there may be some preferential if not obvious TCRβ characteristics that make them better fit.

**Figure 8:**
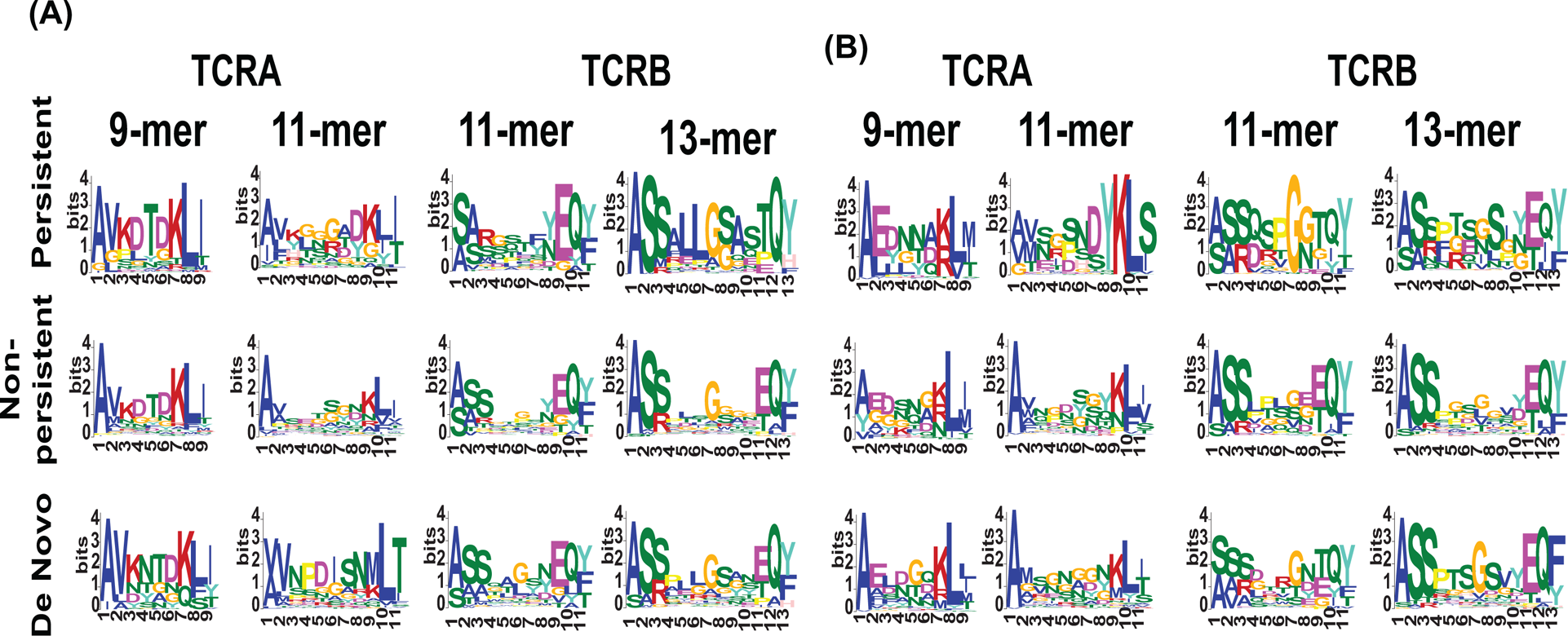
In both YVL-BR and GLC-BM responses, persistent clonotypes have characteristic CDR3α and β motifs that are distinct from non-persistent clonotypes. The *de novo* clonotypes appear to have new and unique CDR3 motifs. HLA-A2/YVL-BR-(A) and GLC-BM-(B) specific TCRα and TCRβ repertoires were analyzed for 3 AIM donors (E1603, E1632, E1655) during the acute (within two weeks of onset of symptoms; primary response) and convalescent (6 months later; memory response) phase of EBV infection. CDR3 motif analysis for the clonotypes within the 3 different subsets, persistent, non-persistent and de novo are shown.

For *de novo* clonotypes, new selection factors appeared that may relate to either a decrease in antigen expression or a change in antigen-expressing cells over the course of persistent infection. For instance, in the YVL-BR 9-mer *de novo* clonotypes, the selection factor AV8.1-AJ34 was maintained in 2/3 donors and a new modified motif, VKNTDK was identified (**Fig S3Ai, 8A, Table S4C**). The *de novo* 11-mer CDR3α response had increased usage of AV12 in all 3 donors (**Fig S3Aii**). In *de novo* BV clonotypes, the pattern of BV-BJ usage changed compared to that observed in AIM.

Similarly, *de novo* 13-mer CDR3β clonotypes were also totally different with usage of a new motif, SALLGX, in 2/3 donors (**Table S4F**).

### GLC-BM persistent, non-persistent and de novo clonotypes have unique characteristics

The persistent GLC-BM TCRα clonotypes maintained the major selection criteria that were identified in AIM with the 9-mer EDNNA motif, which strongly associated with AV5-1-AJ31, being present in a mean 5±3.7% or 10±8.6% of all persistent clonotypes during AIM or CONV, respectively, in all 3 donors (**Fig 8B, Table S4G**). The fact that clonotypes using this motif were not present in non-persistent clonotypes suggests that this motif, and not just the gene family, may be important in determining persistence of GLC-BM-specific clonotypes. The persistent GLC-BM-repertoire also maintained the major selection criteria that were identified in AIM, with the 11-mer SARD motif that strongly associated with BV20.1-BJ1 being present in a mean 16±9.9% or 24±13.7% of all persistent clonotypes during AIM or CONV, respectively in all 3 donors. Two of the donors had the 11-mer SQSPGG motif (**Table S4I**) in a mean 40±8% and 30±25% of all persistent clonotypes during AIM or CONV, respectively.

Only the SARD motif clonotypes appeared in non-persistent BV clonotypes during AIM but at a lower mean frequency of 3±1% (**Table S4J**). The *de novo* clonotype selection appeared to be driven by different factors than the persistent. Although there was much greater diversity and more variation between patients in *de novo* clonotypes (each donor is private) with recruitment of private AV families such as AV41 or AV24 in E1632 and E1655, there was still a preferential usage by 2/3 donors of AV5.1 (**Fig S5i**) and the appearance in 2/3 donors of a new 11-mer CDR3α motif “ELDGQ”, which associated with AV5.1-AJ16.1 (**Fig 8B, Table S4H**). *De novo* clonotypes were also diverse and private using uncommon BV like BV7, BV3 but also using common BV families such as BV20 (**Fig S6**) expressing the SARD motif in 5%±2.9 of *de novo* clonotypes (**Fig 8B, Table S4K**).

In conclusion, the persistent clonotypes made up the vast majority of the AIM and CONV responses. For the most part, the non-persistent clonotypes did not have a motif despite the observation that some of them used a public TCRα or β; this suggests that one of the strongest selection factors for persistence was the CDR3 motif. Additionally, the fact that persistent clonotypes retained features that were identified in AIM further supports their validity. Altogether, these results suggest that the HLA-A2-YVL-BR- or GLC-BM-specific structure contributes strongly to the selection of dominant persistent clonotypes.

## Discussion

This is the first study to use deep sequencing to comprehensively investigate the TCRα and β repertoires to two different EBV epitope-specific CD8 T cell responses over the course of primary infection. We show that while epitope-specific TCR repertoires are highly diverse and vary greatly between donors, they are dominated by distinct clonotypes with public features that persist into convalescence. These persistent clonotypes have distinct features specific to each antigen that appear to drive their peripheral selection; they account for only 9% of unique clonotypes, but predominate in acute infection and convalescence, accounting for 57%±4 of the total epitope-specific response. Surprisingly, the majority of highly diverse unique clonotypes were not detected following AIM and are replaced in convalescence by equally diverse “*de-novo*” clonotypes (43% ± 5% of the total response).

The deep sequencing results show a highly diverse TCR repertoire in each epitope-specific response with 1,292-15,448 and 1,644-7,631 unique clonotypes detected within the YVL-BR and GLC-BM-specific TCR-repertoires, respectively. Such diversity has been underappreciated for the GLC-BM-specific TCR repertoire, with prior studies reporting an oligoclonal repertoire (52, 53, 55). Despite this enormous diversity, there was considerable bias. Although the TCR repertoire was individualized (i.e., each donor studied had a unique TCR-repertoire), there was prevalent and public usage of particular TCRV families such as AV8 within the YVL-BR-specific responses and AV5, AV12 and BV14, BV20 within the GLC-BM-specific populations.

One mechanism which may lead to the dominant public usage and persistence of these clonotypes is that they have TCR features that increase their probability of generation, i.e. they are potentially easier to derive. One of these features, convergent recombination in both the TCRα as well as the TCRβ CDR3 region appears to play a major role in the selection of these persistent clonotypes for expansion and maintenance into long-term memory. This is evidenced by persistent clonotypes using more amino acids that have multiple ways of being derived. A second feature is the usage of shorter germline-derived CDR3 regions with fewer nucleotide additions. The selection of unique public TCR repertoire features, such as CDR3 length, particular TCRAV or BV family usage and motifs, for each epitope in clonotypes that dominate and persist suggest that these clones may be the best fit TCR to recognize the pertinent pMHC complex. In contrast, the broad repertoire of unique clonotypes that are activated in AIM, which is marked by a high viral load and increased inflammation, may not fit as well and perhaps do not receive a TCR signal that leads to survival into memory. Interestingly, 6 months after the initial infection, a completely new (*de novo)* and similarly diverse TCR repertoire has expanded. Continued antigenic exposure in persistent EBV infection may contribute to the evolution of the TCR repertoire overtime.

Prior studies using similar techniques to study influenza A virus (IAV) (not a persistent virus) HLA-A2-restricted IAV-M1_58-67_ and cytomegalovirus (CMV)-pp65 epitope-specific memory responses showed a similar focused diversity of epitope-specific TCR repertoires, suggesting that this is a general principle of antigen-specific repertoire structure (29, 30). Altogether, these studies suggest that the pMHC structure drives selection of the particular public featured dominant clonotypes for each epitope. The broad fluctuating private repertoires show the resilience of memory repertoires and may lend plasticity to antigen recognition, perhaps assisting in early cross-reactive CD8 T cell responses to heterologous new pathogens (28, 56, 57) while at the same time potentially protecting against T cell clonal loss and viral escape (58).

It is, however, possible that this difference in the private diverse portion of the epitope-specific TCR repertoire between acute and convalescence may result from sampling error as we are not able to analyze the full blood volume of an individual. In order to at least partially address this we have analyzed TCRAV and BV deep sequencing data from tetramer-sorted influenza A-M1_58_-specific CD8 T cells (not a persistent virus, thus not influencing TCR repertoire evolution) from one healthy donor of a similar age from two time points one year apart. We compared the TCR overlap of this antigen-specific population at two time points to the donors with AIM in the manuscript. We calculated the overlap between clonotypes at two distinct visits (v1 vs. v7) using the Jaccard similarity coefficient *J*, which is defined as the size of the intersection divided by the size of the union of two sets of clonotypes *A* and B. The mean Jaccard similarity coefficient for TCRAV including both EBV epitopes during AIM was 0.075±0.01 (n=6) and for TCRBV was 0.075±0.01 (n=6). A higher Jaccard similarity coefficient was observed in the healthy donor for TCRVA (0.172) and for TCRVB (0.208). The much higher Jaccard coefficients obtained for the healthy donor suggest that the low overlap between clonotypes observed for acute vs. convalescent visits in EBV infected individuals would not be due to sampling alone. Also, the significant differences in the characteristics of the TCR repertoires of the non-persistent and *de novo* populations would suggest that these are different populations.

There have been limited reports of the importance of TCRα in viral epitope-specific responses. Biased TRAV12.2 usage with CDR1α interaction with the MHC has been observed with the HLA-A2-restricted yellow fever virus epitope, LLWWNGPMAV (59). HLA-B*35:08 restricted EBV BZLF1-specific responses appear to be biased in both TCRα and TCRβ usage, much like HLA-A2-restricted EBV-BR, (60, 61) with a strong preservation of a public TCRα clonotype, AV19-CALSGFYNTDKLIF-J34, which can pair with a few different TCRβ chains. TCRα chain motifs have also been described for HLA-A2-restricted influenza A M1**_58-67_** (IAV-M1), but these appear to make minor contributions to the pMHC-TCR interaction, which is almost completely dominated by CDR3β (29, 45, 46).

The TCR repertoire of the HLA-A2-restricted IAV-M1 epitope is highly biased towards the *TRBV19* gene usage in many individuals and displays a strong preservation of a dominant xRSx CDR3β motif. Crystal structures of TCR specific to this epitope have revealed that the TCR is β-centric with the conserved arginine in the CDR3β loop being inserted into a pocket formed between the peptide and the α2-helix of the HLA-A2 (29, 62). The TCRα has little role in pMHC engagement and this helps explain the high degree of the variability in the CDR3α of sequence and conservation in the CDR3β region. Similarly, previous studies using EBV-GLC-BM-specific CD8 T cells have documented that TCR-pMHC binding modes also contribute to TCR biases (63). The highly public HLA-A2-restricted EBV-GLC-BM-specific AS01 TCR, is highly selected because of a few very strong interactions of its TRAV5- and TRBV20-encoded CDR3 loops with the peptide/MHC.

The present TCR deep sequencing studies, thus reinforce our previous report of an under-appreciated role for TCRα-driven selection of the EBV-YVL-BR-specific repertoire (Fig 6) (51). To the best of our knowledge, our combined studies are among the first to describe a TCR CDR3α-driven selection of viral epitope-specific TCRs with minimal contribution by the TCRBV. The AV8.1 family was used by all individuals and dominated the conserved 9-mer response; it obligately paired with AJ34, and had a predominant CDR3 motif “VKDTDK”, representing 42% and 19% of the total persistent response in AIM and CONV, respectively. In contrast, the BV response was highly diverse without evidence of a strong selection factor, suggesting that AV8.1-VKDTDK-AJ34 could pair with multiple different BV and still successfully be selected by YVL-BR-MHC. In contrast, we did not find any of these AV8.1-VKDTDK-AJ34 expressing TCR in a survey deep sequencing of sorted naïve phenotype CD45RA+, CCR7+ CD8 T cells from 3 age-matched, healthy individuals (one EBV serologically negative and two EBV serologically positive). These results suggest that this clonotype is not inherently present at a high frequency in the naive repertoire, but requires interaction with EBV-YVL-BR to be selected and expanded to these high frequencies.

In contrast, the selection of EBV-GLC-BM-specific TCR repertoire was driven by strong interactions with both chains of TCR, α and β, such as AV5.1-EDNNA-AJ31, BV14-SQSPGG-BJ2 and BV20.1-SARD-BJ1, previously identified public features (43, 52, 53, 55). In a recent study comparing TCRα and β repertoires of various human and murine viral epitopes, none of the responses were primarily driven by interaction with TCRα alone; rather they were predominantly driven by strong interactions with TCRβ or a combination of TCRα and β (11). This apparent preference of YVL-BR TCR repertoires for particular TCRα may create a large repertoire of different memory TCRβ that could potentially cross-react with other ligands such as IAV-M1**_58_**, which predominantly interact with TCRβ (11, 27, 29).

Using single-cell paired TCRαβ sequencing of tetramer sorted CD8 T cells *ex vivo*, we have previously reported that at the at the clonal level recognition of the HLA-A2-restricted EBV-YVL-BR epitope is mainly driven by the TCRα chain (51). The CDR3α motif, KDTDKL, resulted from an obligate AV8.1-AJ34 pairing. This observation coupled with the fact that this public AV8.1-KDTDKL-AJ34 TCR pairs with multiple different TCRβ chains within the same donor (median 4; range: 1-9), suggests that there are some unique structural features of the interaction between the YVL-BR/MHC and the AV8.1-KDTDKL-AJ34 TCR that leads to this high level of selection. TCR motif algorithms identified a lysine at position 1 of the CDR3α motif that is highly conserved and likely important for antigen recognition. Crystal structure analysis of the YVL-BR/HLA-A2 complex revealed that the MHC-bound peptide bulges at position 4, exposing a negatively charged aspartic acid that may interact with the positively charged lysine of CDR3α. TCR cloning and site-directed mutagenesis of the CDR3α lysine ablated EBV-BR-tetramer staining and function. Interestingly, we had previously used TCR structural modeling of the EBV-YVL-BR/MHC complex to predict the occurrence of this important protuberant lysine which might impact TCR interaction (64). Future structural analyses would be important to ascertain whether the YVL-BR TCRα contributes the majority of contacts with the pMHC.

Altogether, our data provide several insights into potential mechanisms of TCR selection and persistence. First, prior studies have revealed that selective use of particular gene families can be explained in part by the fact that the specificity of TCR for a pMHC complex is determined by contacts made between the germline-encoded regions within a V segment and the MHC (63, 65). We show here a highly unique observation of a viral epitope-specific response being strongly selected based not only on a particular TCRAV usage but a highly dominant CDR3α motif and AV-AJ pairing (i.e., the YVL-BR-specific AV8.1-VKDTDK-AJ34 clonotype), with very little role for the TCRBV. Second, it has been suggested that public TCR represent clonotypes present at high frequency in the naïve precursor pool as they may be easier to generate in part as a result of bias in the recombination machinery (66) or convergent recombination of key contact sites (35, 37, 43, 63). Our data demonstrate that convergent recombination of TCRα, as well as TCRβ, may play a dominant role in peripheral selection of clonotypes that persistently detected through memory. As previously reported for TCRβ (35, 37, 43, 63), public clonotypes had a greater probability of being generated.

They used more convergent amino acids than private clonotypes, not only in the CDR3β, but also in the CDR3α. YVL-BR TCRβ, which interestingly is not a strong selection factor for persistent clonotypes did not have public clonotypes with features that led to greater probablity of being generated. Finally, we have previously reported that TCR immunodominance patterns also seem to scale with number of specific interactions required between pMHC and TCR (29). It would seem that TCR that find simpler solutions to being generated and to recognizing antigen are easier to evolve and come to dominate the memory pool (29). Consistent with this our data demonstrate that the dominant persistent clonotypes used shorter predominantly germline derived CDR3α.

Despite the apparent non-persistence of the vast majority of the initial pool of clones deployed during acute infection, clonotypic diversity remained high in memory as a result of the recruitment of a diverse pool of new clonotypes. In a murine model, adoptive transfer of epitope-specific CD8 T cells of known BV families from a single virus-infected mouse to a naive mouse, followed by viral challenge, resulted in altered hierarchy of the clonotypes and the recruitment of new clonotypes, thus maintaining diversity (67). A highly diverse repertoire should allow resilience against loss of individual clonotypes with aging (45) and against skewing of the response after infection with a cross-reactive pathogen (68–71). The large number of clonotypes contributes to the overall memory T cell pool, enhancing the opportunity for protective heterologous immunity now recognized to be an important aspect of immune maturation (56, 72, 73). A large pool of TCR clonotypes could also provide increased resistance to viral escape mutants common in persistent virus infections (58). Finally, different TCR may activate antigen-specific cell functions differently, leading to a more functionally heterogeneous pool of memory cells (74).

In summary, our data reveal that apparent molecular constraints are associated with TCR selection and persistence in the context primary EBV infection. They also show that TCR CDR3α alone can play an equally important role to CDR3β in TCR selection and persistence of important immunodominant responses. Thus, to understand the rules of TCR selection, both TCRα and TCRβ repertoires should be studied. Such studies could elucidate which of the features of the epitope-specific CD8 TCR are associated with an effective response and control of EBV replication or disease.

## Materials and Methods Study population

Three individuals of the age of 18 (E1603, E1632, E1655) who presented with clinical symptoms consistent with acute infectious mononucleosis (AIM) and laboratory studies indicative of primary infection (positive serum heterophile antibody and EBV viral capsid antigen (VCA)-specific IgM) were studied as described (27). Blood samples were collected in heparinized tubes at clinical presentation with AIM symptoms (acute phase) and six months later (memory phase). PBMC were extracted by Ficoll-Paque density gradient media.

### Ethics Statement

The Institutional Review Board of the University of Massachusetts Medical School approved these studies (IRB protocol #: H-3698). All human subjects were adult and provided written informed consent.

### Flow cytometry and isolation of YVL-BR- and GLC-BM-specific CD8 T cells

The percentages of peripheral blood antigen-specific CD8 T cells were measured using flow cytometry analysis. Antibodies included: anti-CD3-FITC, anti-CD4-AF700 and anti-CD8-BV786, 7AAD and PE-conjugated HLA-A*02:01-peptide tetramers (BRLF-1_109-117_: **<u>YVL</u>**DHLIVV; BMLF-1_280-288_: **<u>GLC</u>**TLVAML). Tetramers were made and underwent quality assurance, as previously described(75). Total CD8 T cells were enriched from PBMC by positive selection using MACS technology (Miltenyi Biotec, Auburn, CA) according to the manufacturer’s protocol. The cells were then stained with anti-CD3, anti-CD4, anti-CD8, 7AAD, and GLC-BM- or YVL-BR-tetramers. Live CD3+, CD8+, and GLC-BM- or YVL-BR-tetramer+ cells were sorted by flow cytometry with achieved >95% of purity (FACSAria III, BD) and were subjected for TCR analysis.

### Analysis of TCRα and β CDR3 regions using deep sequencing

The total RNA isolated from minimum 10,000 tetramer+ CD8 T cells was reversely transcribed into cDNA and sent to Adaptive Biotechnologies for TCRα and β-chain profiling following the protocols and standards for sequencing and error correction that comprise ImmunoSEQ platform. In summary, PCR amplification of the CDR3 region is performed using specialized primers that anneal to the V and J recombination regions. Unique molecular identifiers are added during library preparation to track template numbers. After sequencing, CDR3 nucleotide regions are identified and clonal copy numbers are corrected for sequencing and PCR error based on known error rates and clonal frequencies. Sequences of CDR3 regions were identified according to the definition founded by the International ImMunoGeneTics collaboration. Deep sequencing data of TCRα and β repertoires were analyzed using ImmunoSEQ Analyzer versions: 2.0 and 3.0, which were provided by Adaptive Biotechnologies. Only productively (without stop codon) rearranged TCRα and TCRβ sequences were used for repertoire analyses, including sequence aa composition and gene-frequency analyses. The frequencies of *AV-AJ* and *BV-BJ* gene combinations were analyzed with subprograms of the ImmunoSEQ Analyzer software and further processed by Microsoft Excel.

#### Circos plots and motif analysis

The V and J gene segment combinations were illustrated as circos plots(76) across different CDR3 aa sequence lengths. Motif analysis was performed using the Multiple EM for Motif Elicitation (MEME) framework(77). Consensus motifs were acquired across different CDR3 lengths and statistics on those motifs were computed with an in-house program called motifSearch and available at http://github.com/thecodingdoc/motifSearch.

### EBV DNA quantitation in B cells

B cells were purified from whole blood using the RosetteSep human B-cell enrichment cocktail according to the manufacturer’s recommendations (StemCell Technologies, Vancouver BC, Canada). Cellular DNA was extracted using QIAGEN DNeasy Blood & Tissue Kit (Valencia, CA). Each DNA sample was diluted to 5ng/ul and the Roche LightCycler EBV Quantitation Kit (Roche Diagnostics, Indianapolis, IN) was used to quantify EBV DNA copy number in the samples as recommended by the manufacturer. Reactions were run in duplicate. B cell counts in each sample were determined using a previously described PCR assay to quantify the copy number of the gene encoding CCR5 (two copies per diploid cell)(78). Samples were normalized to B cell counts and EBV DNA copy number was calculated as DNA copy per 10^6^ B cells.

### Convergence Analyses

The number of unique nucleotide sequences encoding an amino acid sequence of TCRAV and TCRBV regions specific for YVL-BR and GLC-BM epitopes were calculated across the pooled repertoires of all individuals. The number of nucleotide additions required to produce a TCRAV or TCRBV sequence was determined by aligning the germline V gene at the 5’ end of the TCRAV or TCRBV sequence and then the J gene segment at the 3’ end of the TCR sequence. The germline D genes were subsequently aligned with nucleotides in the junction between the identified V and J regions. Nucleotides identified in the junctions between the V, D, and J gene segments were considered to be nucleotide additions. The significance values are based on multivariant two-way ANOVA.

## Statistics

GraphPad Prism version 7.0 for Mac OSX (GraphPad Software, La Jolla, CA) was used for all statistical analyses.

## Data availability

Raw TCR deep sequencing data can be accessed: https://urldefense.proofpoint.com/v2/url?u=http-3Aclients.adaptivebiotech.com&d=DwQGaQ&c=WJBj9sUF1mbpVIAf3biu3CPHX4MeRjY_w4DerPlOmhQ&r=p6IL5ohbVyB2IGgNCmdbh-A5IMFqxKtq0WBpidjH1QE&m=hueuAoY7ZXzP9YMFmhGPKpu9iLorr5nv05XTqQklDuI&s=AdlhcrGwYqZ-QWYlQON5AJFRO88HSQe1qPUMaWRkQik&e=

The login information is as follows:

Username: ">

Password: gil-2018-review

## Abbreviations

EBV: Epstein Barr virus
TCR: T cell receptor
AIM: acute infectious mononucleosis
pMHC: peptide major histocompatibility complex
CDR3: complementarity determining region 3
CONV: convalescence
YVL-BR: BRLF-1_109_ epitope
GLC-BM: BMLF-1_280_ epitope.

## Acknowledgements

We are grateful to the study subjects for their participation; to George Corey and Jessica Conrad for obtaining clinical samples; Linda Lambrecht, Robin Brody, Anita Gautam, and Jennifer Henderson for expert technical assistance; Margaret McManus for data management and critical review of the manuscript; and Dr. Raymond Welsh for critical review of the manuscript. This research used resources of the Advanced Photon Source, a U.S. Department of Energy (DOE) Office of Science User Facility operated for the DOE Office of Science by Argonne National Laboratory under Contract No. DE-AC02-06CH11357. Use of the Lilly Research Laboratories Collaborative Access Team (LRL-CAT) beamline at Sector 31 of the Advanced Photon Source was provided by Eli Lilly Company, which operates the facility.

## Author contributions

L.K.S. and K.L. obtained samples and conceived the study. A.G. and L.K. contributed to study design, and were primarily responsible for cell sorting and TCR sequencing. All authors contributed to data analyses. D.G. and R.C. performed all computational analyses. L.K.S., K.L., L.K., A.G. and D.G. assumed primary responsibility for writing the manuscript. All authors reviewed, provided substantive input, and approved of the final manuscript.

## Supporting Information Legends

**Figure S1: 9-mer TCRα AV8.1-VKDTDK-AJ34 drives the selection of YVL-specific CD8 T-cells in AIM.** The TCR repertoire is deconstructed by analyzing V family usage in pie chart format, CDR3 length analyses, VJ pairing by using circos plot analyses, and CDR3 amino acid motif analyses using Multiple MEME framework(77). **(i)** YVL-BR-specific TCRVA (A) and TRVB (B) repertoires are analyzed for 3 AIM donors (E1603, E1632, E1655) during AIM. Frequency of each TRAV (A) and TRBV (B) in total YVL-BR-specific TCR-repertoire is shown in pie charts (i). The pie plots are labeled with gene families having a frequency ≥5%. The total numbers of unique clonotypes in each donor is shown below the pie charts. (ii) CDR3 length distribution along with (iii) circos plots depicting V-J gene pairing and (iiii) CDR3 motif analysis for the clonotypes with the two most dominant CDR3 lengths. Genes are colored by V gene family with a fixed color sequence used throughout the manuscript.

**Figure S2: TCRα, AV5-EDNNA-AJ31, and TCRβ, BV14-SQSPGG-BJ2 and BV20-SARD-BJ1, clones are dominant selection factors for GLC-BM-specific CD8 T-cells during AIM.** GLC-BM-specific TCRVA (A) and TRVB (B) repertoires are analyzed for 3 AIM donors (E1603, E1632, E1655) during AIM. Frequency of each TRAV (A) and TRBV (B) in total GLC-BM-specific TCR-repertoire is shown in pie charts (i). The pie plots are labeled with gene families having a frequency ≥5%. The total numbers of unique clonotypes in each donor is shown below the pie charts. There is consistent usage of AV5 and AV12 genes in all 3 donors. There is consistent usage of BV20 in all 3 donors. Otherwise there is a high degree of variability in other AV and BV usage between donors. (ii) CDR3 length distribution along with (iii) circos plots depicting V-J gene pairing and (iiii) motif analysis for the clonotypes with the two most dominant CDR3 lengths. The frequencies of V-J combinations are displayed in circos plots, with frequency of each V or J cassette represented by its arc length and that of the V-J cassette combination by the width of the arc.

**Figure S3: Unique patterns of V-J usage for persistent, non-persistent and de novo clonotypes 9-mer (i) and 11-mer (ii) CDR3α of the YVL-BR-specific CD8 T cell responses as obtained by deep sequencing.** The frequencies of V-J combinations in three AIM donors for YVL-BR-specific TCRα repertoires are displayed in circos plots, with frequency of each V or J cassette represented by its arc length and that of the V-J cassette combination by the width of the arc. For comparison the total acute and convalescence circus plots are also shown.

**Figure S4: Unique patterns of V-J usage for persistent, non-persistent and de novo clonotypes of 11-mer (i) and 13-mer (ii) CDR3β of the YVL-specific CD8+ T cell responses as obtained by deep sequencing.** The frequencies of V-J combinations in three AIM donors for GLC-BM-specific TCRβ repertoires are displayed in circos plots, with frequency of each V or J cassette represented by its arc length and that of the V-J cassette combination by the width of the arc.

**Figure S5: Unique patterns of V-J usage for persistent, non-persistent and *de novo* clonotypes of 9-mer (i) and 11-mer (ii) CDR3α of the GLC-BM-specific CD8 T-cell responses.** The frequencies of V-J combinations in three AIM donors for GLC-BM-specific TCRαβ repertoires are displayed in circos plots, with frequency of each V or J cassette represented by its arc length and that of the V-J cassette combination by the width of the arc. For comparison the total acute and convalescence circos plots are also shown.

**Figure S6: Unique patterns of V-J usage for persistent, non-persistent and *de novo* clonotypes of 11-mer (i) and 12-mer (ii) CDR3β of the GLC-BM-specific CD8 T-cell responses.** The frequencies of V-J combinations in three AIM donors for GLC-BM-specific TCRαβ repertoires are displayed in circos plots, with frequency of each V or J cassette represented by its arc length and that of the V-J cassette combination by the width of the arc.

**Supplemental Table 1.** Characteristics of AIM donors in study population.

**Supplemental Table S2.** TCR Sequencing depth and counts of productive DNA rearrangements by donor, epitope-specificity and time point.

**Supplemental Table S3:** YVL-BR specific and GLC-BM specific TRAV and TRBV dominant motifs.

**Supplemental Table S4:** YVL-BR specific and GLC-BM specific TRAV and TRBV dominant motifs in persistent, non-persistent and *de novo* clonotypes.

## References

1. Taylor GS, Long HM, Brooks JM, Rickinson AB, Hislop AD. 2015. The immunology of Epstein-Barr virus-induced disease. Annu Rev Immunol 33:787–821.

2. Luzuriaga K, Sullivan JL. 2010. Infectious mononucleosis. N Engl J Med 362:1993–2000.

3. Henle G, Henle W, Clifford P, Diehl V, Kafuko GW, Kirya BG, Klein G, Morrow RH, Munube GM, Pike P, Tukei PM, Ziegler JL. 1969. Antibodies to Epstein-Barr virus in Burkitt’s lymphoma and control groups. J Natl Cancer Inst 43:1147–1157.

4. Pender MP, Burrows SR. 2014. Epstein-Barr virus and multiple sclerosis: potential opportunities for immunotherapy. Clin Transl Immunology 3:e27.

5. Crawford DH. 2001. Biology and disease associations of Epstein-Barr virus. Philos Trans R Soc Lond B Biol Sci 356:461–473.

6. Loren AW, Porter DL, Stadtmauer EA, Tsai DE. 2003. Post-transplant lymphoproliferative disorder: a review. Bone Marrow Transplant 31:145–155.

7. Bollard CM, Gottschalk S, Torrano V, Diouf O, Ku S, Hazrat Y, Carrum G, Ramos C, Fayad L, Shpall EJ, Pro B, Liu H, Wu MF, Lee D, Sheehan AM, Zu Y, Gee AP, Brenner MK, Heslop HE, Rooney CM. 2014. Sustained complete responses in patients with lymphoma receiving autologous cytotoxic T lymphocytes targeting Epstein-Barr virus latent membrane proteins. J Clin Oncol 32:798–808.

8. Bollard CM, Rooney CM, Heslop HE. 2012. T-cell therapy in the treatment of post-transplant lymphoproliferative disease. Nat Rev Clin Oncol 9:510–519.

9. Pender MP, Csurhes PA, Burrows JM, Burrows SR. 2017. Defective T-cell control of Epstein-Barr virus infection in multiple sclerosis. Clin Transl Immunology 6:e126.

10. Keymeulen B, Vandemeulebroucke E, Ziegler AG, Mathieu C, Kaufman L, Hale G, Gorus F, Goldman M, Walter M, Candon S, Schandene L, Crenier L, De Block C, Seigneurin JM, De Pauw P, Pierard D, Weets I, Rebello P, Bird P, Berrie E, Frewin M, Waldmann H, Bach JF, Pipeleers D, Chatenoud L. 2005. Insulin needs after CD3-antibody therapy in new-onset type 1 diabetes. N Engl J Med 352:2598–2608.

11. Dash P, Fiore-Gartland AJ, Hertz T, Wang GC, Sharma S, Souquette A, Crawford JC, Clemens EB, Nguyen THO, Kedzierska K, La Gruta NL, Bradley P, Thomas PG. 2017. Quantifiable predictive features define epitope-specific T cell receptor repertoires. Nature 547:89–93.

12. Glanville J, Huang H, Nau A, Hatton O, Wagar LE, Rubelt F, Ji X, Han A, Krams SM, Pettus C, Haas N, Arlehamn CSL, Sette A, Boyd SD, Scriba TJ, Martinez OM, Davis MM. 2017. Identifying specificity groups in the T cell receptor repertoire. Nature 547:94–98.

13. DeWitt WS, Smith A, Schoch G, Hansen JA, Matsen FA, Bradley P. 2018. Human T cell receptor occurrence patterns encode immune history, genetic background, and receptor specificity. Elife 7.

14. Emerson RO, DeWitt WS, Vignali M, Gravley J, Hu JK, Osborne EJ, Desmarais C, Klinger M, Carlson CS, Hansen JA, Rieder M, Robins HS. 2017. Immunosequencing identifies signatures of cytomegalovirus exposure history and HLA-mediated effects on the T cell repertoire. Nat Genet 49:659–665.

15. Attaf M, Huseby E, Sewell AK. 2015. alphabeta T cell receptors as predictors of health and disease. Cell Mol Immunol 12:391–399.

16. Attaf M, Sewell AK. 2016. Disease etiology and diagnosis by TCR repertoire analysis goes viral. Eur J Immunol 46:2516–2519.

17. Cohen JI, Mocarski ES, Raab-Traub N, Corey L, Nabel GJ. 2013. The need and challenges for development of an Epstein-Barr virus vaccine. Vaccine 31 Suppl 2:B194–196.

18. Yanagi Y, Yoshikai Y, Leggett K, Clark SP, Aleksander I, Mak TW. 1984. A human T cell-specific cDNA clone encodes a protein having extensive homology to immunoglobulin chains. Nature 308:145–149.

19. Zinkernagel RM, Doherty PC. 1974. Restriction of in vitro T cell-mediated cytotoxicity in lymphocytic choriomeningitis within a syngeneic or semiallogeneic system. Nature 248:701–702.

20. Hedrick SM, Cohen DI, Nielsen EA, Davis MM. 1984. Isolation of cDNA clones encoding T cell-specific membrane-associated proteins. Nature 308:149–153.

21. La Gruta NL, Gras S, Daley SR, Thomas PG, Rossjohn J. 2018. Understanding the drivers of MHC restriction of T cell receptors. Nat Rev Immunol 18:467–478.

22. Siu G, Clark SP, Yoshikai Y, Malissen M, Yanagi Y, Strauss E, Mak TW, Hood L. 1984. The human T cell antigen receptor is encoded by variable, diversity, and joining gene segments that rearrange to generate a complete V gene. Cell 37:393–401.

23. Miles JJ, Douek DC, Price DA. 2011. Bias in the alphabeta T-cell repertoire: implications for disease pathogenesis and vaccination. Immunol Cell Biol 89:375–387.

24. Nikolich-Zugich J, Slifka MK, Messaoudi I. 2004. The many important facets of T-cell repertoire diversity. Nat Rev Immunol 4:123–132.

25. Furman D, Jojic V, Sharma S, Shen-Orr SS, Angel CJ, Onengut-Gumuscu S, Kidd BA, Maecker HT, Concannon P, Dekker CL, Thomas PG, Davis MM. 2015. Cytomegalovirus infection enhances the immune response to influenza. Sci Transl Med 7:281ra243.

26. Kloverpris HN, McGregor R, McLaren JE, Ladell K, Harndahl M, Stryhn A, Carlson JM, Koofhethile C, Gerritsen B, Kesmir C, Chen F, Riddell L, Luzzi G, Leslie A, Walker BD, Ndung’u T, Buus S, Price DA, Goulder PJ. 2015. CD8+ TCR Bias and Immunodominance in HIV-1 Infection. J Immunol 194:5329–5345.

27. Aslan N, Watkin LB, Gil A, Mishra R, Clark FG, Welsh RM, Ghersi D, Luzuriaga K, Selin LK. 2017. Severity of Acute Infectious Mononucleosis Correlates with Cross-Reactive Influenza CD8 T-Cell Receptor Repertoires. MBio 8.

28. Watkin LB, Gil A, Mishra R, Aslan N, Ghersi D, Luzuriaga K, Selin LK. 2017. Potential of influenza A memory CD8+ T-cells to protect against Epstein Barr virus (EBV) seroconversion. Journal of Allergy and Clinical Immunology 140:1206–1210.

29. Song I, Gil A, Mishra R, Ghersi D, Selin LK, Stern LJ. 2017. Broad TCR repertoire and diverse structural solutions for recognition of an immunodominant CD8+ T cell epitope. Nat Struct Mol Biol 24:395–406.

30. Chen G, Yang X, Ko A, Sun X, Gao M, Zhang Y, Shi A, Mariuzza RA, Weng NP. 2017. Sequence and Structural Analyses Reveal Distinct and Highly Diverse Human CD8+ TCR Repertoires to Immunodominant Viral Antigens. Cell Rep 19:569–583.

31. Abdel-Hakeem MS, Boisvert M, Bruneau J, Soudeyns H, Shoukry NH. 2017. Selective expansion of high functional avidity memory CD8 T cell clonotypes during hepatitis C virus reinfection and clearance. PLoS Pathog 13:e1006191.

32. Klarenbeek PL, Remmerswaal EB, ten Berge IJ, Doorenspleet ME, van Schaik BD, Esveldt RE, Koch SD, ten Brinke A, van Kampen AH, Bemelman FJ, Tak PP, Baas F, de Vries N, van Lier RA. 2012. Deep sequencing of antiviral T-cell responses to HCMV and EBV in humans reveals a stable repertoire that is maintained for many years. PLoS Pathog 8:e1002889.

33. Sant S, Grzelak L, Wang Z, Pizzolla A, Koutsakos M, Crowe J, Loudovaris T, Mannering SI, Westall GP, Wakim LM, Rossjohn J, Gras S, Richards M, Xu J, Thomas PG, Loh L, Nguyen THO, Kedzierska K. 2018. Single-Cell Approach to Influenza-Specific CD8(+) T Cell Receptor Repertoires Across Different Age Groups, Tissues, and Following Influenza Virus Infection. Front Immunol 9:1453.

34. Welsh RM, Kim SK, Cornberg M, Clute SC, Selin LK, Naumov YN. 2006. The privacy of T cell memory to viruses. Curr Top Microbiol Immunol 311:117–153.

35. Venturi V, Kedzierska K, Price DA, Doherty PC, Douek DC, Turner SJ, Davenport MP. 2006. Sharing of T cell receptors in antigen-specific responses is driven by convergent recombination. Proc Natl Acad Sci U S A 103:18691–18696.

36. Venturi V, Chin HY, Price DA, Douek DC, Davenport MP. 2008. The role of production frequency in the sharing of simian immunodeficiency virus-specific CD8+ TCRs between macaques. J Immunol 181:2597–2609.

37. Venturi V, Price DA, Douek DC, Davenport MP. 2008. The molecular basis for public T-cell responses? Nat Rev Immunol 8:231–238.

38. Fazilleau N, Cabaniols JP, Lemaitre F, Motta I, Kourilsky P, Kanellopoulos JM. 2005. Valpha and Vbeta public repertoires are highly conserved in terminal deoxynucleotidyl transferase-deficient mice. J Immunol 174:345–355.

39. Mora TW, A.M. 2016. Quantifying lymphocyte diversity. q-Bio-PE https://arxiv.org/pdf/1604.00487.pdf.

40. Lim A, Trautmann L, Peyrat MA, Couedel C, Davodeau F, Romagne F, Kourilsky P, Bonneville M. 2000. Frequent contribution of T cell clonotypes with public TCR features to the chronic response against a dominant EBV-derived epitope: application to direct detection of their molecular imprint on the human peripheral T cell repertoire. J Immunol 165:2001–2011.

41. Argaet VP, Schmidt CW, Burrows SR, Silins SL, Kurilla MG, Doolan DL, Suhrbier A, Moss DJ, Kieff E, Sculley TB, Misko IS. 1994. Dominant selection of an invariant T cell antigen receptor in response to persistent infection by Epstein-Barr virus. J Exp Med 180:2335–2340.

42. Sethna Z, Elhanati Y, Callan CG, Walczak AM, Mora T. 2019. OLGA: fast computation of generation probabilities of B- and T-cell receptor amino acid sequences and motifs. Bioinformatics 35:2974–2981.

43. Venturi V, Chin HY, Asher TE, Ladell K, Scheinberg P, Bornstein E, van Bockel D, Kelleher AD, Douek DC, Price DA, Davenport MP. 2008. TCR beta-chain sharing in human CD8+ T cell responses to cytomegalovirus and EBV. J Immunol 181:7853–7862.

44. Venturi V, Quigley MF, Greenaway HY, Ng PC, Ende ZS, McIntosh T, Asher TE, Almeida JR, Levy S, Price DA, Davenport MP, Douek DC. 2011. A mechanism for TCR sharing between T cell subsets and individuals revealed by pyrosequencing. J Immunol 186:4285–4294.

45. Gil A, Yassai MB, Naumov YN, Selin LK. 2015. Narrowing of human influenza A virus-specific T cell receptor alpha and beta repertoires with increasing age. J Virol 89:4102–4116.

46. Naumov YN, Naumova EN, Yassai MB, Kota K, Welsh RM, Selin LK. 2008. Multiple glycines in TCR alpha-chains determine clonally diverse nature of human T cell memory to influenza A virus. J Immunol 181:7407–7419.

47. Yang X, Gao M, Chen G, Pierce BG, Lu J, Weng NP, Mariuzza RA. 2015. Structural Basis for Clonal Diversity of the Public T Cell Response to a Dominant Human Cytomegalovirus Epitope. J Biol Chem 290:29106–29119.

48. Pymm P, Illing PT, Ramarathinam SH, O’Connor GM, Hughes VA, Hitchen C, Price DA, Ho BK, McVicar DW, Brooks AG, Purcell AW, Rossjohn J, Vivian JP. 2017. MHC-I peptides get out of the groove and enable a novel mechanism of HIV-1 escape. Nat Struct Mol Biol 24:387–394.

49. Yang X, Chen G, Weng NP, Mariuzza RA. 2017. Structural basis for clonal diversity of the human T-cell response to a dominant influenza virus epitope. J Biol Chem 292:18618–18627.

50. Singh NK, Riley TP, Baker SCB, Borrman T, Weng Z, Baker BM. 2017. Emerging Concepts in TCR Specificity: Rationalizing and (Maybe) Predicting Outcomes. J Immunol 199:2203–2213.

51. Kamga L, Gil A, Song I, Brody R, Ghersi D, Aslan N, Stern LJ, Selin LK, Luzuriaga K. 2019. CDR3alpha drives selection of the immunodominant Epstein Barr virus (EBV) BRLF1-specific CD8 T cell receptor repertoire in primary infection. PLoS Pathog 15:e1008122.

52. Annels NE, Callan MF, Tan L, Rickinson AB. 2000. Changing patterns of dominant TCR usage with maturation of an EBV-specific cytotoxic T cell response. J Immunol 165:4831–4841.

53. Callan MF, Fazou C, Yang H, Rostron T, Poon K, Hatton C, McMichael AJ. 2000. CD8(+) T-cell selection, function, and death in the primary immune response in vivo. J Clin Invest 106:1251–1261.

54. Nguyen TH, Bird NL, Grant EJ, Miles JJ, Thomas PG, Kotsimbos TC, Mifsud NA, Kedzierska K. 2017. Maintenance of the EBV-specific CD8+ TCRalphabeta repertoire in immunosuppressed lung transplant recipients. Immunol Cell Biol 95:77–86.

55. Levitsky V, de Campos-Lima PO, Frisan T, Masucci MG. 1998. The clonal composition of a peptide-specific oligoclonal CTL repertoire selected in response to persistent EBV infection is stable over time. J Immunol 161:594–601.

56. Cornberg M, Clute SC, Watkin LB, Saccoccio FM, Kim SK, Naumov YN, Brehm MA, Aslan N, Welsh RM, Selin LK. 2010. CD8 T cell cross-reactivity networks mediate heterologous immunity in human EBV and murine vaccinia virus infections. J Immunol 184:2825–2838.

57. Welsh RM, Selin LK. 2002. No one is naive: the significance of heterologous T-cell immunity. Nat Rev Immunol 2:417–426.

58. Wolfl M, Rutebemberwa A, Mosbruger T, Mao Q, Li HM, Netski D, Ray SC, Pardoll D, Sidney J, Sette A, Allen T, Kuntzen T, Kavanagh DG, Kuball J, Greenberg PD, Cox AL. 2008. Hepatitis C virus immune escape via exploitation of a hole in the T cell repertoire. J Immunol 181:6435–6446.

59. Bovay A, Zoete V, Dolton G, Bulek AM, Cole DK, Rizkallah PJ, Fuller A, Beck K, Michielin O, Speiser DE, Sewell AK, Fuertes Marraco SA. 2018. T cell receptor alpha variable 12-2 bias in the immunodominant response to Yellow fever virus. Eur J Immunol 48:258–272.

60. Tynan FE, Burrows SR, Buckle AM, Clements CS, Borg NA, Miles JJ, Beddoe T, Whisstock JC, Wilce MC, Silins SL, Burrows JM, Kjer-Nielsen L, Kostenko L, Purcell AW, McCluskey J, Rossjohn J. 2005. T cell receptor recognition of a ‘super-bulged’ major histocompatibility complex class I-bound peptide. Nat Immunol 6:1114–1122.

61. Tynan FE, Borg NA, Miles JJ, Beddoe T, El-Hassen D, Silins SL, van Zuylen WJ, Purcell AW, Kjer-Nielsen L, McCluskey J, Burrows SR, Rossjohn J. 2005. High resolution structures of highly bulged viral epitopes bound to major histocompatibility complex class I. Implications for T-cell receptor engagement and T-cell immunodominance. J Biol Chem 280:23900–23909.

62. Ishizuka J, Stewart-Jones GB, van der Merwe A, Bell JI, McMichael AJ, Jones EY. 2008. The structural dynamics and energetics of an immunodominant T cell receptor are programmed by its Vbeta domain. Immunity 28:171–182.

63. Miles JJ, Bulek AM, Cole DK, Gostick E, Schauenburg AJ, Dolton G, Venturi V, Davenport MP, Tan MP, Burrows SR, Wooldridge L, Price DA, Rizkallah PJ, Sewell AK. 2010. Genetic and structural basis for selection of a ubiquitous T cell receptor deployed in Epstein-Barr virus infection. PLoS Pathog 6:e1001198.

64. Antunes DA, Rigo MM, Freitas MV, Mendes MFA, Sinigaglia M, Lizee G, Kavraki LE, Selin LK, Cornberg M, Vieira GF. 2017. Interpreting T-Cell Cross-reactivity through Structure: Implications for TCR-Based Cancer Immunotherapy. Front Immunol 8:1210.

65. Davis MM, McHeyzer-Williams M, Chien YH. 1995. T-cell receptor V-region usage and antigen specificity. The cytochrome c model system. Ann N Y Acad Sci 756:1–11.

66. Yassai M, Bosenko D, Unruh M, Zacharias G, Reed E, Demos W, Ferrante A, Gorski J. 2011. Naive T cell repertoire skewing in HLA-A2 individuals by a specialized rearrangement mechanism results in public memory clonotypes. J Immunol 186:2970–2977.

67. Cukalac T, Chadderton J, Handel A, Doherty PC, Turner SJ, Thomas PG, La Gruta NL. 2014. Reproducible selection of high avidity CD8+ T-cell clones following secondary acute virus infection. Proc Natl Acad Sci U S A 111:1485–1490.

68. Selin LK, Brehm MA, Naumov YN, Cornberg M, Kim SK, Clute SC, Welsh RM. 2006. Memory of mice and men: CD8+ T-cell cross-reactivity and heterologous immunity. Immunol Rev 211:164–181.

69. Selin LK, Wlodarczyk MF, Kraft AR, Nie S, Kenney LL, Puzone R, Celada F. 2011. Heterologous immunity: immunopathology, autoimmunity and protection during viral infections. Autoimmunity 44:328–347.

70. Clute SC, Naumov YN, Watkin LB, Aslan N, Sullivan JL, Thorley-Lawson DA, Luzuriaga K, Welsh RM, Puzone R, Celada F, Selin LK. 2010. Broad cross-reactive TCR repertoires recognizing dissimilar epstein-barr and influenza A virus epitopes. J Immunol 185:6753–6764.

71. Clute SC, Watkin LB, Cornberg M, Naumov YN, Sullivan JL, Luzuriaga K, Welsh RM, Selin LK. 2005. Cross-reactive influenza virus-specific CD8+ T cells contribute to lymphoproliferation in Epstein-Barr virus-associated infectious mononucleosis. J Clin Invest 115:3602–3612.

72. Benn CS, Netea MG, Selin LK, Aaby P. 2013. A small jab - a big effect: nonspecific immunomodulation by vaccines. Trends Immunol 34:431–439.

73. Selin LK, Brehm MA. 2007. Frontiers in nephrology: heterologous immunity, T cell cross-reactivity, and alloreactivity. J Am Soc Nephrol 18:2268–2277.

74. Han A, Glanville J, Hansmann L, Davis MM. 2014. Linking T-cell receptor sequence to functional phenotype at the single-cell level. Nat Biotechnol 32:684–692.

75. Luzuriaga K, McManus M, Catalina M, Mayack S, Sharkey M, Stevenson M, Sullivan JL. 2000. Early therapy of vertical human immunodeficiency virus type 1 (HIV-1) infection: control of viral replication and absence of persistent HIV-1-specific immune responses. J Virol 74:6984–6991.

76. Krzywinski M, Schein J, Birol I, Connors J, Gascoyne R, Horsman D, Jones SJ, Marra MA. 2009. Circos: an information aesthetic for comparative genomics. Genome Res 19:1639–1645.

77. Bailey TL, Elkan C. 1994. Fitting a mixture model by expectation maximization to discover motifs in biopolymers. Proc Int Conf Intell Syst Mol Biol 2:28–36.

78. Precopio ML, Sullivan JL, Willard C, Somasundaran M, Luzuriaga K. 2003. Differential kinetics and specificity of EBV-specific CD4+ and CD8+ T cells during primary infection. J Immunol 170:2590–2598.

